# Active Learning Enables Efficient Directed Evolution of a Far-Red Fluorescent Protein with Minimal Experimental Data

**DOI:** 10.64898/2026.08.12.744534

**Authors:** Daniel V. Brown, Ryan S. Cross, Shiying Zhu, Thomas Hill, Chhon Ling Sok, Misty R. Jenkins, Marija Dramicanin, Rory Bowden

## Abstract

Fluorescent proteins are fundamental tools for cellular imaging. Most fluorescent proteins in routine use, including GFP, are derived from the jellyfish *Aequorea victoria* and emit blue-green light, which is strongly absorbed and scattered by tissue, limiting imaging depth. Far-red and near-infrared fluorescent proteins, engineered from bacteriophytochromes, address this limitation because far-red light penetrates tissue considerably further. However, these proteins are typically much dimmer than their *A. victoria* -derived counterparts. Improving brightness by conventional directed evolution requires screening large random mutant libraries, a process that is slow, labor-intensive, and often impractical outside specialized laboratories. We utilized an active-learning-guided directed evolution workflow that identified improved variants from substantially less data than conventional screening. Each round coupled automated, miniaturized cell-free protein expression directly from a DNA template without cloning or cell culture, with a machine-learning model retrained on cumulative sequence–function data to nominate the most informative variants for the next round. Applied to miRFP670nano3, this workflow screened 120 variants across successive rounds and identified twelve with improved brightness, the best four-fold brighter in bacterial systems. However, these gains did not translate when the variants were evaluated in mammalian cells, indicating that performance can be strongly dependent on cellular context. Retrospective simulation across benchmark datasets from ProteinGym showed that performing more experimental batches with fewer samples per batch consistently accelerated convergence to high-fitness sequences. Incorporating protein-language-model derived zero-shot fitness priors also accelerated convergence, but only in proportion to how well each prior score correlated with the true fitness landscape. Together, these findings established generalizable design rules, favoring smaller acquisition batches and confidence-weighted priors, for engineering proteins from minimal experimental data.

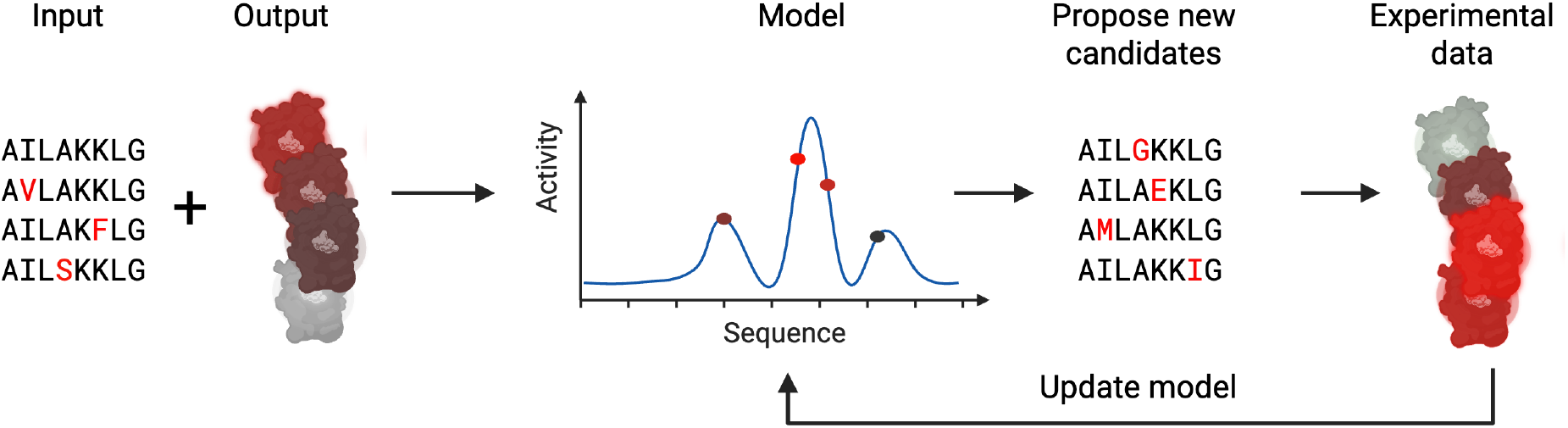

## Introduction

Fluorescent proteins (FPs) have revolutionized cellular and molecular biology, enabling real-time visualization of gene expression, protein localization, and complex cellular processes in living systems. ^1^ Originally derived from the jellyfish *Aequorea victoria*, the green fluorescent protein (GFP) and its engineered derivatives have become ubiquitous tools for imaging and biosensing applications.

Despite their widespread utility, *A. victoria* FPs are limited by several intrinsic properties. Most notably, their chromophore maturation requires oxygen, reducing their signal in hypoxic environments. Additionally, GFP and related proteins are excited by blue or green light, which is subject to significant scattering and absorption in biological tissues. As a result, imaging in whole organisms or deep tissues remains challenging. ^2^ Beta-barrel-based farred protein variants have been developed but also require oxygen. ^3^

To overcome these limitations, far-red and near-infrared FPs have been developed, primarily derived from bacteriophytochromes. Among these, the miRFPnano family are small, monomeric, and exhibit excitation and emission maxima in the far-red region, where light penetrates tissue more effectively and background autofluorescence is minimized. ^4,5^ However, these proteins suffer from relatively low brightness compared to their GFP counterparts.

Directed evolution has proven to be a powerful method for optimizing protein properties, including brightness, stability, and spectral characteristics. ^6^ Traditional directed evolution campaigns rely on generating and screening large libraries of random protein variants to identify improved mutants. While effective, this process is labor-intensive, time-consuming and costly, especially when the screening throughput is limited or when functional assays are complex.

Recent advances in machine learning (ML) offer exciting new possibilities for protein engineering. ^7,8^ By leveraging sequence–function data, ML-guided approaches can efficiently predict and prioritize beneficial mutations, thereby reducing the experimental burden. Active-learning directed evolution (ALDE) closes the loop between ML and experimental data. At each round, the trained model queries the sequences expected to be most informative or highest-performing, and the resulting measurements are fed back to update its predictions before the next round of selection. ^8^

In this study, we present an ALDE campaign to enhance the brightness of miRFP670nano3, a far-red fluorescent protein. ^9^ Using a combination of low-volume, automated laboratory cycles and cell-free expression systems, we rapidly generated and tested miRFP670nano3 protein variants. This strategy enabled us to quickly converge on variants with increased brightness, including mutations that would have been difficult to predict from structural or sequence conservation analyses alone.

## Results

### Active-learning directed evolution improves miRFP670nano3 brightness

To improve the brightness of miRFP670nano3, we established an ALDE workflow (Fig 1a). We evaluated surrogate models on nine different deep mutational scanning (DMS) datasets of varying difficulty. No significant difference in final-best fitness emerged across models (Friedman test, p = 0.32; Fig S1b). Therefore we selected Gaussian process (GP) regression on PCA-reduced protein language model (pLM) embeddings primarily for the posterior uncertainty estimates it provides for Bayesian optimization acquisition, in addition to its slight (non-significant) advantage in median best-variant performance (Fig S1b,c). We then developed a laboratory assay to measure brightness using a cell-free protein expression system (CFPS) with an N-terminal HiBiT tag to measure protein expression. Throughout our study, “brightness” refers to fluorescence signal normalized to the parental sequence, which is distinct from molecular brightness. Coupled with laboratory automation at nanoliter volumes, this assay showed sensitivity to input DNA template concentration (Fig S1d). We conducted five rounds of ALDE: rounds 1–3 measured single mutants by weighted activity, and those single mutants whose brightness or expression exceeded parental miRFP670nano3 were subsequently combined in all possible 3-way combinations for rounds 4 and 5. 74 of 120 variants tested differed significantly in weighted activity from parental miRFP670nano3 (one-sample t-test, BH-adjusted p < 0.05), including 12 with more than a two-fold increase in weighted activity and 8 with more than a two-fold decrease (Fig 1b). The top variant K6G-L50F-A143T achieved 4.35-fold higher brightness at the cost of 25% reduced expression. ALDE worked as intended with a progressively higher average weighted activity across experimental rounds (Fig 1c).

**Figure 1:**
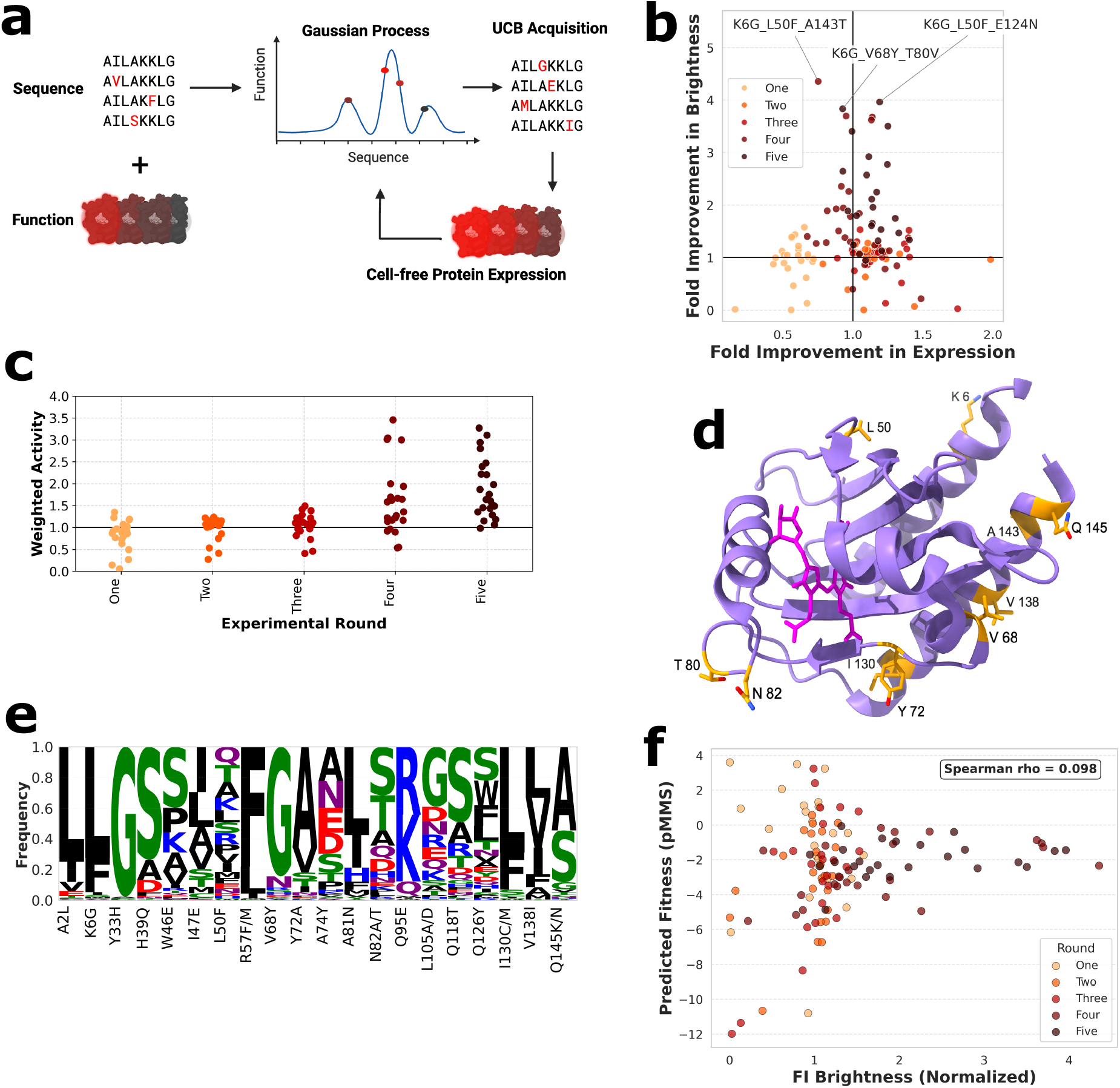
Active learning and CFPS improve miRFP670nano3 brightness. (a) Schematic of the active-learning directed workflow. (b) Brightness and expression of miRFP670nano3 variants across five rounds of ALDE. The top three variants are labeled. (c) Weighted activity (75% brightness, 25% expression) across five rounds of ALDE. (d) miRFP670nano3 structure (purple) overlaid with weighted-activity-promoting mutations (gold) and biliverdin (pink). (e) Sequence logo of natural residue frequencies at the positions nominated by ALDE, computed across the 1000 GAF domains most similar to miRFP670nano3. (f) pLM-predicted fitness (pMMS) vs relative brightness measured by CFPS.

Overlaying the top mutations suggested by our model onto the miRFP670nano3 experimental structure (PDB: 7LSC) revealed no obvious mechanistic basis for brightness improvement (Fig 1d). Notably, mutations proximal to the biliverdin binding site were consistently detrimental. This includes C86A, which disrupts the cysteine that forms the covalent thioether bond to the biliverdin chromophore. We examined the residue suggestion frequencies and observed the model increasingly focused on certain positions, repeatedly selecting K6 (as K6G) and exploring multiple alternate residues at Y72 (Fig S1e).

We next checked if the mutations suggested by the model were novel or already existing in nature, given the ALDE input features were pLM embeddings. pLM embeddings have been demonstrated to contain information similar to a multiple sequence alignment and capture evolutionary relationships. ^10^ The mutations conferring higher brightness were in a mixture of both categories (Fig 1e). In some cases such as A2L, the mutation reverted to the most common amino acid seen across evolution. In other cases such as K6G the mutation was novel. Novel mutations were preferentially obtained from sequence positions with lower Shannon entropy (Two-sided Mann-Whitney U test, p = 0.032; Fig S1f). There was a weak, non-significant positive correlation between fitness predicted by a pLM (predicted masked-marginal score, pMMS) and brightness as measured in our assays (Spearman *ρ* = 0.098, *p* = 0.286, n = 120; Fig 1f), which is consistent with other ALDE campaigns. ^8^ Repeated 5-fold cross-validation showed no classic ML model, including a Gaussian process, generalized well to held-out variants, indicating model class was not the limiting factor (Fig S1g).

### Biological validation of engineered miRFP670nano3 variants

After improving miRFP670nano3 brightness by ALDE and CFPS we checked whether the results validate in more complex expression systems. We transfected the parental sequence and three distinct high-performing variants from the CFPS campaign into HEK293T cells (Fig S2a,b). In contrast to CFPS, in mammalian cells the parental sequence had the highest far-red geometric mean fluorescence intensity (geoMFI), 3.6-to 4.1-fold higher than the three engineered variants (one-way ANOVA, *p* = 2.0*×* 10^*−*6^; Welch’s post-hoc t-tests all significant; Fig 2a,b). Surprisingly, the GFP geoMFI obtained from the P2A-linked eGFP construct was lowest for the parental sequence suggesting lower protein abundance in transfected cells (Fig 2b). However live cell counts were not reduced for parental sequence relative to the variants (Fig 2b), arguing against a toxicity-driven effect. We also confirmed higher fluorescent intensity of the parental sequence in a construct without P2A-linked GFP (Fig. S2c).

**Figure 2:**
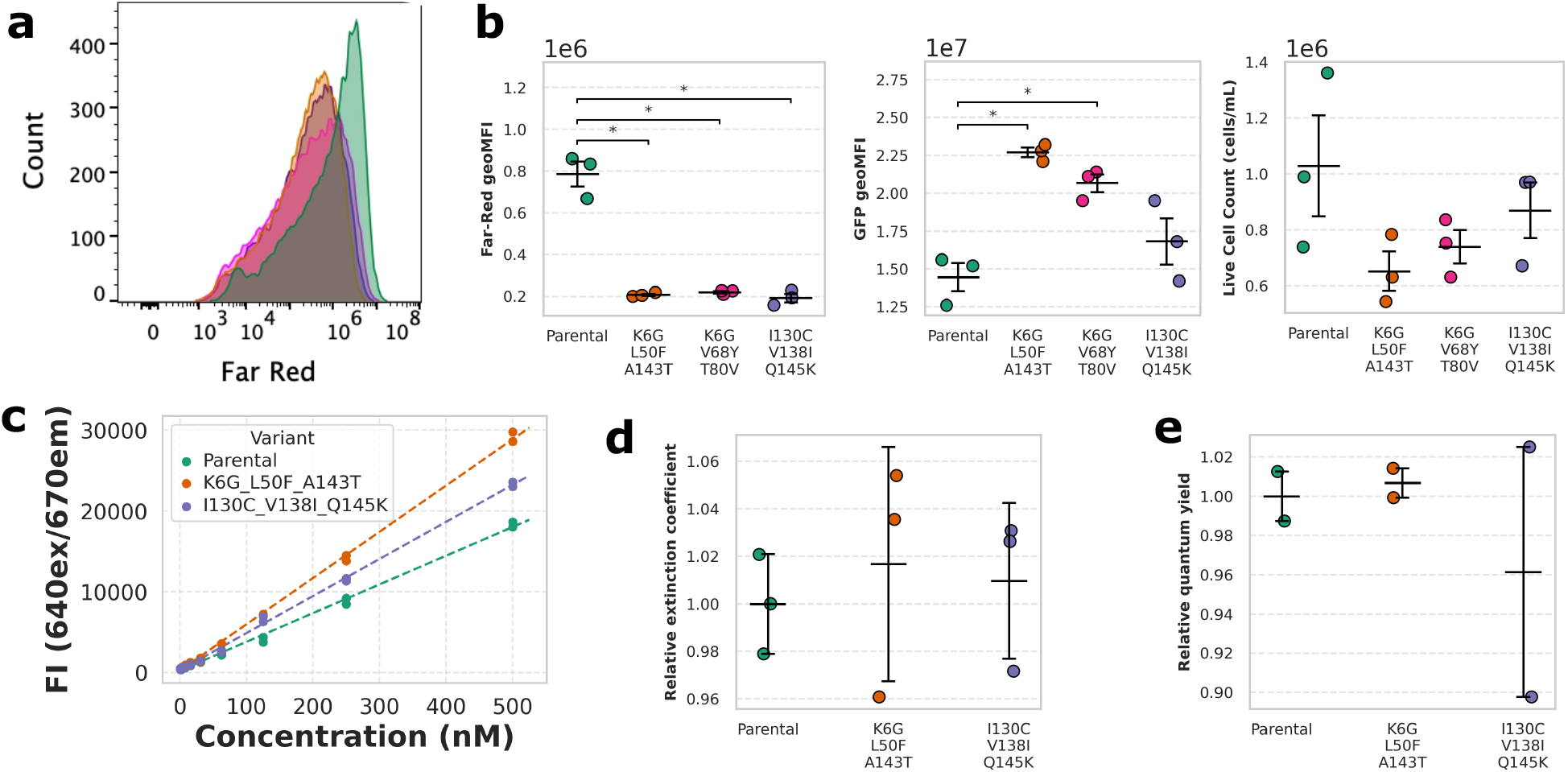
Biological characterization of engineered miRFP670nano3 variants. (a) Far-red fluorescence histograms of HEK293T cells transiently transfected with parental miRFP670nano3 or engineered variants. (b) Far-red geometric mean fluorescence intensity (left), GFP geometric mean fluorescence intensity (middle), and live cell count (right) for parental and engineered variants. (c) Fluorescence intensity of purified parental and engineered variant proteins. (d) Relative extinction coefficient of purified parental and engineered variants. (e) Relative quantum yield measured by the gradient method and normalized to the parental sequence.

We additionally made stable miRFP670nano3 expressing cell lines via lentiviral transduction and conducted titration experiments which confirmed the parental sequence was brighter in mammalian cells (Fig. S2d). Fluorescence fell off more steeply with decreasing viral dose for the parental sequence compared to K6G-L50F-A143T (14% steeper, BH-adjusted pairwise *p* = 0.003; Fig. S2e). Given the ALDE campaign was conducted in a purified bacterially derived CFPS system we next checked performance of miRFP670nano3 variants in bacteria. Following purification from bacterial lysates, K6G-L50F-A143T and I130C-V138I-Q145K variants were brighter than the parental sequence (one-way ANOVA *p* = 0.0011, pairwise contrasts all significant, *p≤* 0.05; Fig 2c). We also confirmed increased brightness of K6G-L50F-A143T in an alternative bacterial lysate-based CFPS system (Fig. S3a). Interestingly both engineered variants were less thermostable than the parental sequence (Fig S3b). Neither excitation, emission, absorbance, extinction coefficient or relative quantum yield properties were altered in any of the variants (Fig 2d,e and Fig S3c,d,e). Spectral and biophysical properties are summarized in Table 1.

**Table 1:** Spectral and biophysical properties of parental miRFP670nano3 and derived variants.

| Variant | $\lambda_{ex}$ (nm) | $\lambda_{em}$ (nm) | $\varepsilon_{rel}$ | $QY_{rel}$ | $T_i$ ($^{\circ}\text{C}$) | Brightness<sub>CFPS</sub> | Brightness<sub>purified</sub> | Brightness<sub>HEK293T</sub> |
| --- | --- | --- | --- | --- | --- | --- | --- | --- |
| Parental (WT) | 648 | 672 | 1.00 | 1.00 | 61.6 | 1.00 | 1.00 | 1.00 |
| K6G-L50F-A143T | 646 | 672 | 1.02 | 1.01 | 52.8 | 4.35 | 1.61 | 0.26 |
| I130C-V138I-Q145K | 647 | 673 | 1.01 | 0.96 | 52.4 | 3.50 | 1.29 | 0.25 |
| K6G-V68Y-T80V | ND | ND | ND | ND | ND | 3.83 | ND | 0.28 |

### Benchmarking ALDE design choices by simulation

Two design choices shape an ALDE campaign before any experimental work begins: how many variants to test per batch, and how to prioritize the first round in the absence of data. We addressed both using retrospective simulation on deep mutational scanning datasets. Within the nine DMS datasets selected in our simulation, CreiLOV is the most similar to miRFP670nano3. Although unrelated in sequence, both proteins are small, monomeric fluorescent proteins that have been engineered to improve brightness for biological imaging applications. ^11^

To check whether the number of variants selected per experimental batch in our ALDE campaign was well chosen, we swept batch size at a fixed budget of 384 total evaluations across the nine benchmark datasets, from a single batch of 384 (random sampling) down to 47 batches of 8 samples. Performance improved as batch size shrank for CreiLOV cumulative-mean fitness (Fig 3a; Fig S4a). This finding extended to the majority of other datasets (mean within-dataset Spearman *ρ* =*−* 0.927, one-sample Wilcoxon signed-rank test vs. zero, *p* = 0.0039; Fig 3b). The effect of reducing batch size was less pronounced though still significant for final best fitness (mean within-dataset Spearman *ρ* = −0.524, *p* = 0.0195; Fig S4b,c).

**Figure 3:**
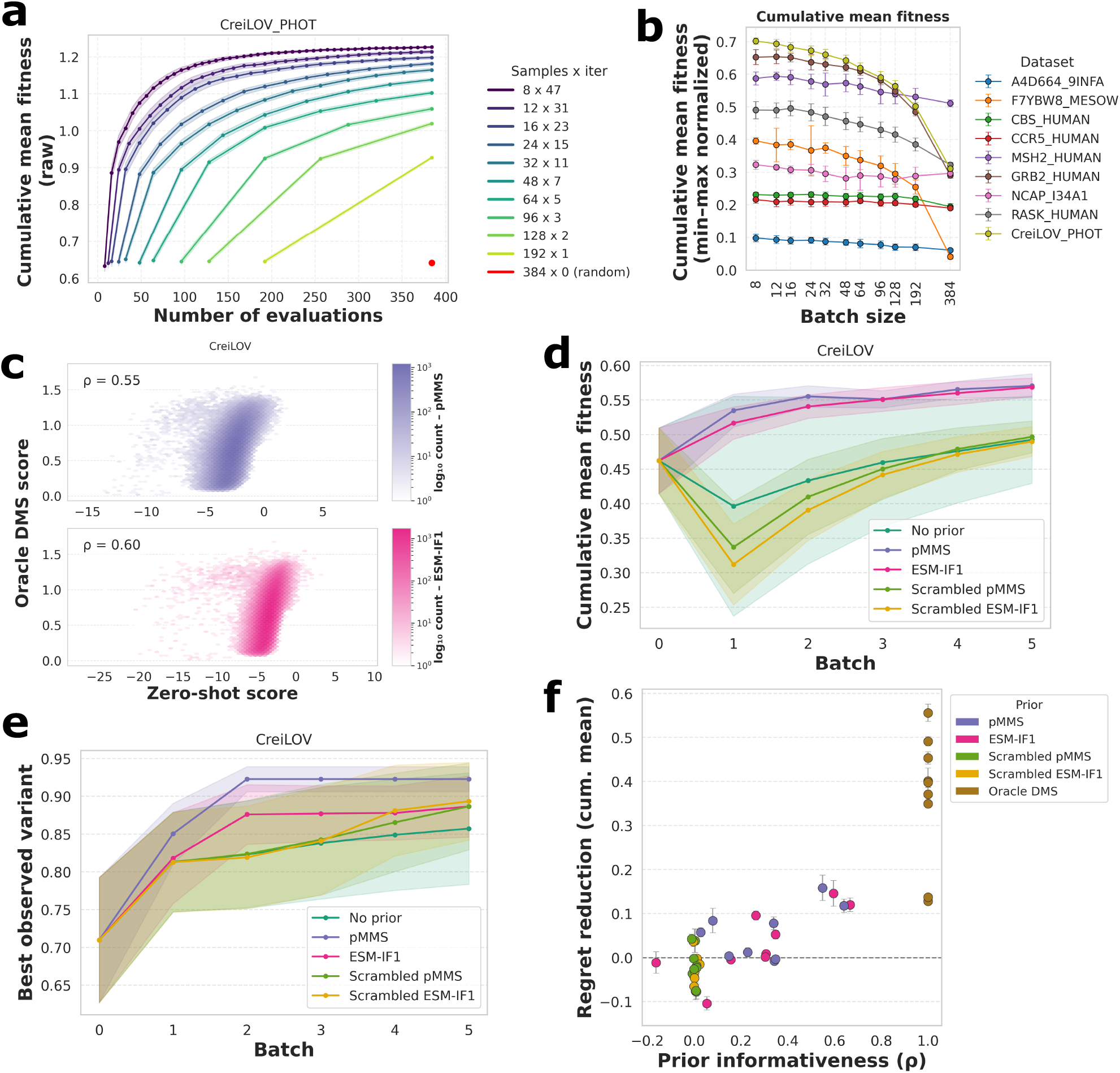
Retrospective simulation improves ALDE experimental design. (a) Cumulative-mean convergence (raw fitness) for CreiLOV across different combinations of batches and samples per batch. Error bars/shading represent sd of 20 random seeds throughout. (b) Cumulative-mean fitness summary (min–max normalized per dataset) across acquisition batch sizes for all nine benchmark datasets. (c) Zero-shot fitness scores versus ground truth DMS fitness for the CreiLOV dataset. Top = predicted masked-marginal score (pMMS). Bottom = ESM-IF1 inverse-folding log-likelihood. (d) Cumulative-mean convergence for the CreiLOV dataset across GP prior means and their scrambled controls. (e) Convergence to the best variant for the CreiLOV dataset across GP prior means. (f) Prior-mean informativeness (Spearman correlation with oracle DMS score) versus regret reduction (improvement in cumulative-mean fitness to the no-prior baseline). The oracle anchor point (*ρ* = 1 by construction) is excluded from statistical analysis.

The first round of an ALDE campaign requires selecting candidate sequences with no experimental data to guide it. We tested whether this cold start could be improved using the zero-shot fitness prior of Benjamins et al., ^12^ in which a protein language model score sets the GP’s prior mean and a learned weight (*α*) governs how strongly that prior is trusted. We compare two simple priors, ESM-2 masked-marginal scores (pMMS) and ESM-IF1 inverse-folding log-likelihoods with their scrambled negative controls. Both zero-shot scores correlated positively with experimental fitness for CreiLOV (pMMS ρ = −0.549; ESM-IF1 *ρ* = 0.596; *n* = 167,529 genotypes; Fig 3c; Fig S5a,b). For CreiLOV, both priors improved cumulative-mean fitness convergence relative to the no-prior baseline and scrambled controls (paired Wilcoxon test, BH-adjusted *p* = 1.9*×* 10^*−*6^; scrambled controls: BH-adjusted *p >* 0.3, Fig 3d). The difference was more modest for best observed variant (Fig 3e). Across all nine datasets, the benefit of using these priors was more modest and strongly protein-dependent (Fig S5c,d). We defined this benefit as the fractional reduction in cumulative-mean regret relative to the no-prior baseline. It scaled with how informative the prior was within each dataset (positive in 8/9 datasets, Wilcoxon signed-rank test vs. zero *p* = 0.0078; mean within-dataset Spearman *ρ* = 0.652; Fig 3f). The importance of the prior (*α*) learned by the GP also tracked informativeness, down weighting priors that were poorly correlated with ground truth (Fig S5e).

## Discussion

The primary motivation for this study was to address the limited brightness of miRFP670nano3 by leveraging ALDE. By coupling machine learning with a targeted experimental design strategy, we aimed to efficiently navigate the sequence–function landscape and identify improved variants with minimal experimental cycles. Our approach required screening 120 total variants across five rounds, a dramatic reduction compared to typical deep mutational scanning or random library screening campaigns. The mutations the model selected would not have been prioritized by structural intuition. Nor would conservation have prioritized them, as novel mutations were preferentially drawn from positions of low Shannon entropy. It is in exactly this situation, where structure and conservation offer little guidance, active learning with experimental data is highly suitable.

Biological validation in mammalian cells did not recapitulate the CFPS gains. The PURExpress reaction is supplied with saturating biliverdin (5 µM) in solution and lacks chaperones or proteasome, whereas biliverdin supplied to mammalian cells (25 µM) is poorly membrane-permeable and inhibited in serum. ^4,13^ A variant selected under cofactor excess has little reason to win under co-factor limitation: if brightness in cells depends on the fraction of protein loaded with biliverdin, a variant never selected for efficient loading would underperform once that cofactor becomes limiting. Validating CFPS hits in mammalian cells each round, as was done for the parental protein, ^9^ would have caught this earlier.

In the context of the bacterially produced purified protein, the engineered variants were brighter than the parental sequence. Thermostability was decreased while the extinction coefficient and quantum yield were unchanged. In the bacterial system biliverdin was produced within the cell using heme oxygenase co-transformation. To some extent this mimics the CFPS system of excess biliverdin being available.

Because miRFP670nano3, like other bacteriophytochrome-derived near-infrared fluorescent proteins, acquires biliverdin post-translationally rather than forming its chromophore autocatalytically, brightness measured per unit protein reflects not only the intrinsic extinction coefficient and quantum yield of the mature holoprotein but also the efficiency with which expressed protein is folded and successfully loaded with the cofactor. We therefore favor a model in which the mutations identified by our screen do not alter the photophysics of the bound chromophore itself, but instead increase the fraction of expressed protein that reaches the folded, biliverdin-loaded, fluorescent state.

Our simulations showed smaller, more frequent batches converge faster to a higher cumulative-mean fitness, though this must be weighed against the turnaround time and cost of ordering DNA and running the assay for each batch. 48–96 samples per batch is a reasonable compromise that does not degrade performance much. The effect was weaker for final-best fitness, which ultimately matters most in directed evolution. However, using the cumulative mean captures batch selection and can diagnose systematic failure, as was the case when all of our variants failed to reproduce the CFPS gain in mammalian cells.

The prior-mean result shows an informative zero-shot score can accelerate ALDE, but only on landscapes where that score correlates with true fitness, a property that can only be known in retrospect for a real campaign. In our ALDE campaign pMMS was uninformative for miRFP670nano3 brightness. Weighting the prior by a learned confidence term offers a principled hedge against this uncertainty as zero-shot score prediction continues to improve. ^14^

Several limitations suggest avenues for future work. Mean pooling of pLM embeddings followed by dimension reduction likely discarded valuable sequence information. Our trained model did not generalize to held-out positions. Richer embedding or pooling strategies could retain more functional signal. ^7,15,16^ Our initial search space was restricted to single-site mutagenesis. In contrast, a campaign targeting combinatorial mutagenesis of the binding pocket may better target mutations that enhance biliverdin binding affinity.

Together these results show that active learning can improve a far-red fluorescent protein from few experimental measurements and that biological context is critical in directed evolution campaigns.

## Methods

Full experimental details are provided in Supporting Methods.

### Cell-free protein expression

Variants were ordered as linear synthetic genes (Integrated DNA Technologies) and expressed with the PUR-Express In Vitro Protein Synthesis Kit (NEB E6800L) supplemented with 5 µM biliverdin (Sigma-Aldrich 30891). Brightness was read on a ClarioStar plate reader (BMG Labtech) and expression was measured from an N-terminal

HiBiT tag. Both readouts were normalized to the mean of the parental sequence and combined into a single weighted activity label (75% brightness, 25% expression) used to train the surrogate model.

### Active-learning initialization

The design space was single-site saturation mutagenesis of miRFP670nano3, with the most destabilizing mutations (upper quartile of FoldX-predicted ΔΔ*G*) removed for the initialization round only. ^17^ All variants remained available in subsequent rounds. Sequences were embedded with ESM3, ^18^ mean pooled across positions, ^8^ column-standardized and reduced to their first three principal components as GP models struggle with high dimensionality. ^7^ The initial 24 variants comprised 12 chosen by k-means clustering of the embeddings (*k* = 12, the variant nearest each centroid) and 12 nominated by HotSpot Wizard as tolerant to mutation across evolution. ^19^

### Interpreting trained models

The miRFP670nano3 protein structure (PDB: 7LSC) was used as the basis for modeling. ^20^ GP generalization was evaluated on the single-mutant subset (n = 72) by repeated 5-fold cross-validation. Evolutionary conservation of the mutations nominated during ALDE was assessed against the top 1000 most similar natural GAF-domain homologs.

### Protein expression and characterization

Bacterial expression plasmids were co-transfected with Heme oxygenase 1 from *Synechocystis* (strain ATCC 27184). After bacterial lysis the His-tagged miRFP670nano3 variants were purified with His-Tag purification resin and buffer exchanged into 20 mM Tris-HCl pH 8.0, 300 mM NaCl, 50 µM EDTA, 10% glycerol, 1 mM DTT, 0.01% NP-40 and analyzed by size-exclusion chromatography to confirm protein purity. Inflection temperature (*T*_*i*_) was tested using a Tycho NT.6 (Nanotemper). Fluorescence spectral properties were characterized using a CLARIOstar plate reader.

### Biological validation experiments in mammalian cell lines

Parental miRFP670nano3 and its variants were transiently transfected in HEK293T cells with a P2A-GFP sequence. Stable lines generated by lentiviral transduction. Farred and GFP fluorescence were measured on a NovoCyte Penteon Flow Cytometer (Agilent). Cell counts were performed with a Countess III (Thermo Fisher).

### Retrospective simulation benchmarking

ALDE was simulated on nine deep mutational scanning datasets from ProteinGym to compare surrogate models against random-sampling and ensemble baselines. Acquisition batch size was swept at a fixed budget of 384 evaluations. Seven zero-shot GP prior-mean conditions were compared against scrambled negative controls and an oracle positive control (20 random seeds throughout).

## Data and code availability

The raw and processed data along with analysis code for this manuscript are available from GitHub. Deep mutational scanning data for ALDE simulations was obtained from ProteinGym on 23 March 2026. Pipelines used to process raw DMS data into ALDE results is also available from GitHub.

## Acknowledgements

We acknowledge the WEHI Advanced Genomics Facility, Research Computing, Flow Cytometry Facility and Instrumentation facility for professional and timely service. M.R.J is supported by a NHMRC Investigator Grant L1. R.S.C. is supported by a The Kids Cancer Project Col Reynolds Mid-Career Fellowship. D.V.B is supported by funding to the Advanced Genomics Facility from the Walter and Eliza Hall Institute. We thank Andrew Xu, Justin Bedo, Niall Geoghegan, Pradeep Rajasekhar and Richard Birkenshaw for helpful discussions. Thank you to Kevin Weston for training on Clariostar plus. Experimental design figures were created using Biorender.com.

## Author Contribution

Conceptualization, D.V.B. Methodology, D.V.B, R.S.C, T.H, S.Z, M.D. Data Acquisition, D.V.B, R.S.C, S.Z, C.L.S. Data Analysis, D.V.B, T.H. Investigation, D.V.B. Writing, D.V.B. Funding Acquisition, D.V.B, R.B. Supervision, D.V.B, M.R.J, M.D, R.B.

## Conflict of Interest

The authors report no conflicts of interest.

## Declaration of AI-assisted technologies in the writing process

During the preparation of this work the author used Claude for copy-editing. After using these tools, D.V.B reviewed and edited the content as needed and takes full responsibility for the content of the published article.

## Supporting information

Supporting Information is available with this manuscript.

- DNA and plasmid sequences used in this study (Table S1).
- Additional analysis of the active-learning directed-evolution (ALDE) campaign, including ProteinGym benchmark, synthetic gene design, learned-model residue preferences and sequence conservation (Figure S1).
- Flow cytometry gating strategy and fluorescence analysis of transiently and stably transduced miRFP670nano3 variants (Figure S2).
- Spectroscopic and visual characterization of purified miRFP670nano3 variants (Figure S3).
- Acquisition batch-size benchmarking (Figure S4).
- Gaussian process prior-mean benchmarking (Figure S5).
- Supporting Methods: cell-free protein expression, active-learning statistics, Gaussian process interpretation and sequence-conservation analysis, protein expression and purification, biochemical characterization, mammalian-cell and lentiviral validation, and retrospective simulation benchmarking.

## Supplementary Tables

**Supp Table 1**

List of gene variants and oligonucleotide sequences used in this study.

Table 2: DNA and plasmid sequences used in this study. Available as excel file in Supplementary Data.

## Supplementary Figures

**Supp figure S1**

Related to figure 1. ALDE setup, CFPS assay development and GP model interpretation.

**Supp figure S2**

Related to figure 2. Additional flow cytometry analysis.

**Supp figure S3**

Related to figure 2. Additional characterization of purified miRFP670nano3.

**Supp figure S4**

Related to figure 3. Additional batch-size benchmarking analysis.

**Supp figure S5**

Related to figure 3. Additional information about GP prior-mean benchmarking.

**Figure S1:**
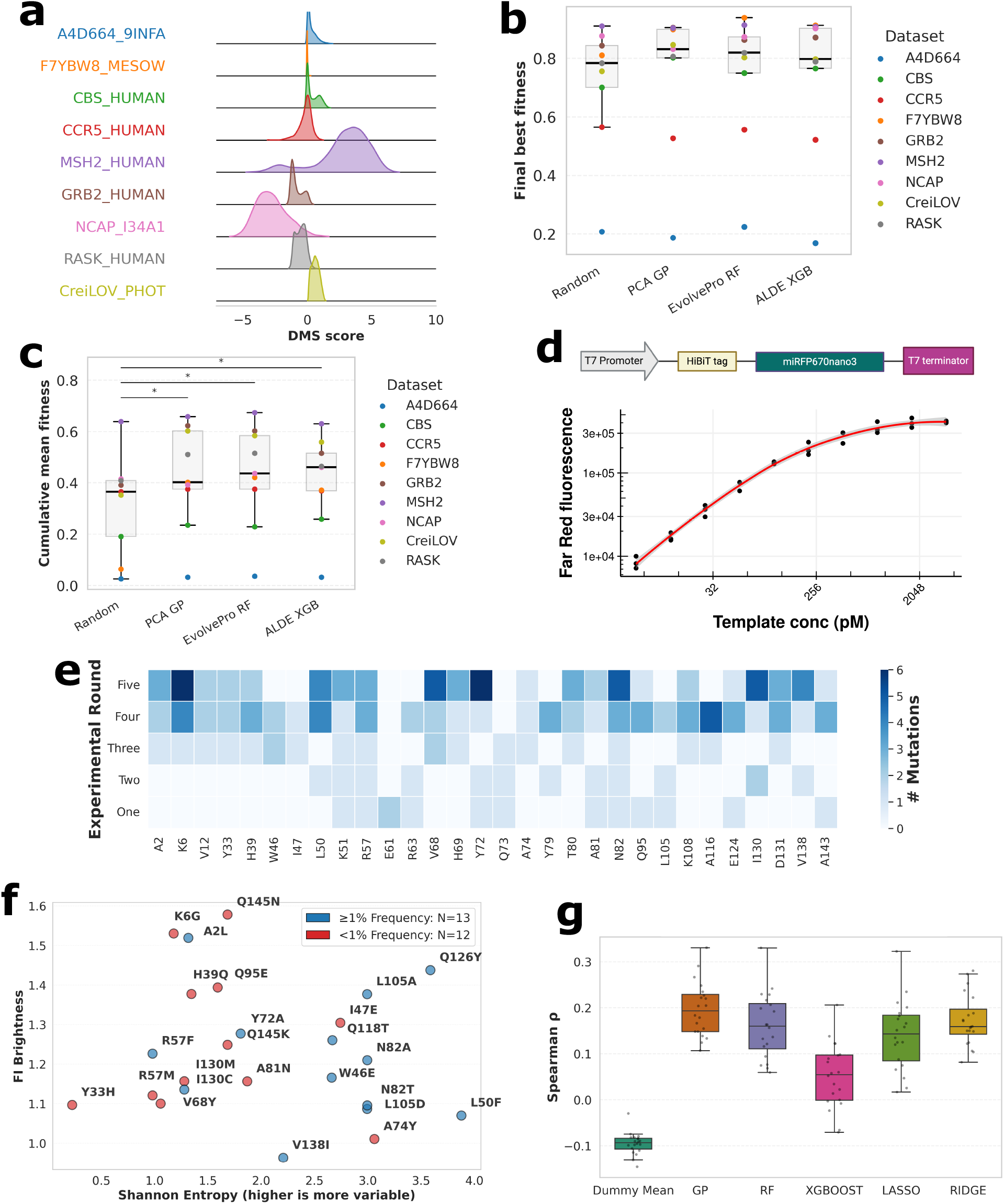
Additional analyses for ALDE campaign. (a) Fitness distributions for nine DMS datasets from ProteinGym. The raw DMS Score is shown on the x-axis (b) Final-best fitness for the nine datasets, comparing three ALDE methods against a random baseline. (c) As for (b), showing cumulative-mean fitness instead. Each point represents the per-protein mean of 20 random seeds. (d) Schematic of the linear synthetic genes used in CFPS assay and dose response of input template. (e) Heatmap of per-residue mutation-recommendation frequency by the surrogate model across the ALDE campaign. (f) Relationship between observed brightness (FI Brightness) and sequence conservation (Shannon entropy) at mutated positions, colored by mutant-residue frequency among natural homologs. (g) GP predictive performance vs. alternative model classes on held-out single-mutant variants (repeated 5-fold CV, 20 random seeds).

**Figure S2:**
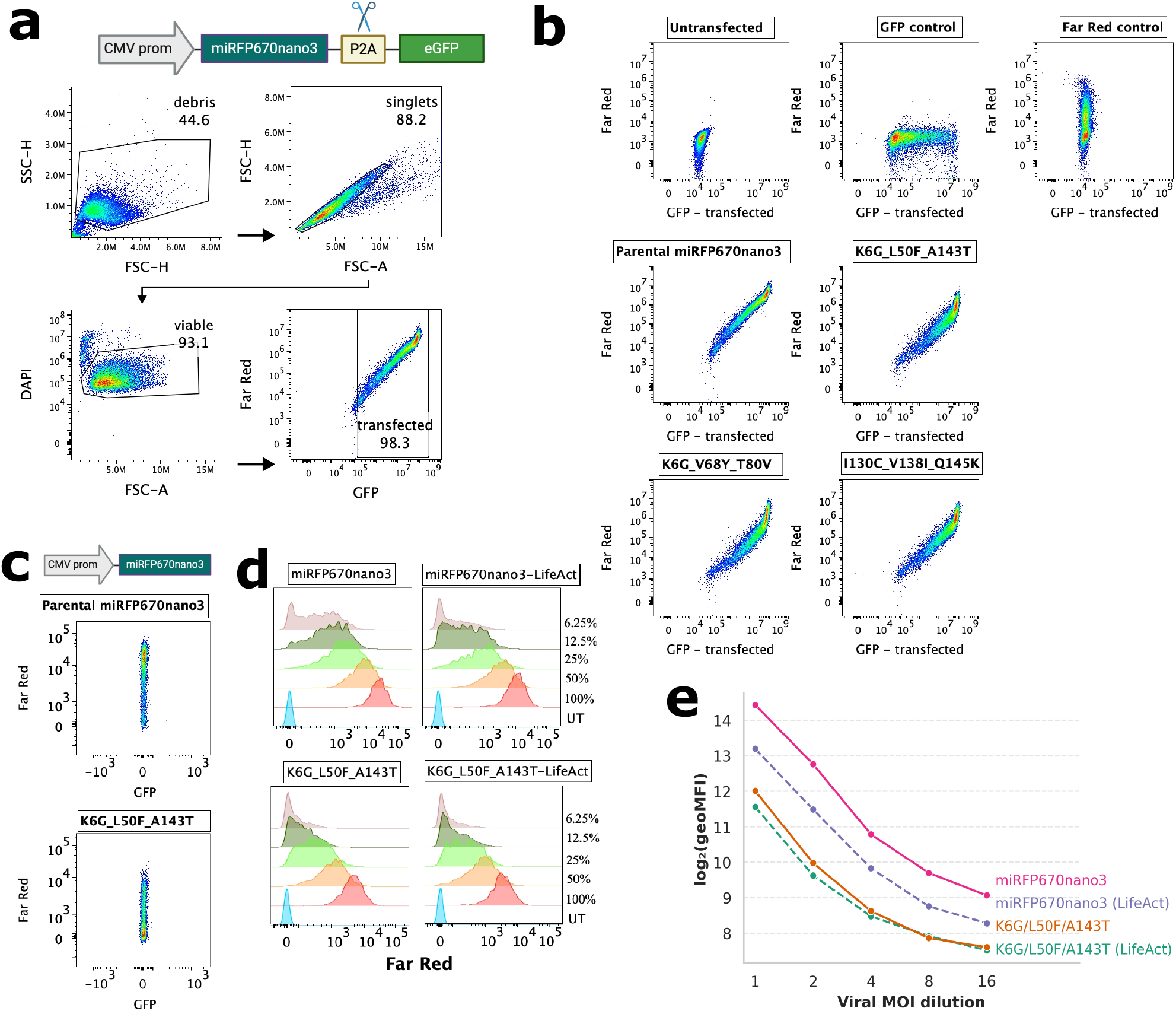
Flow cytometry gating and per-variant scatter. (a) Reporter construct schematic (CMV-miRFP670nano3-P2A-eGFP) and sequential gating hierarchy applied to parental miRFP670nano3. (b) Bivariate scatter plots of miRFP670nano3 far-red versus P2A-linked eGFP fluorescence. (c) Bivariate scatter plots of miRFP670nano3 transfections without P2A-GFP. (d) Serial-dilution far-red fluorescence histograms of stable lentiviral lines. (e) Dose-response of log(geoMFI) versus viral MOI dilution.

**Figure S3:**
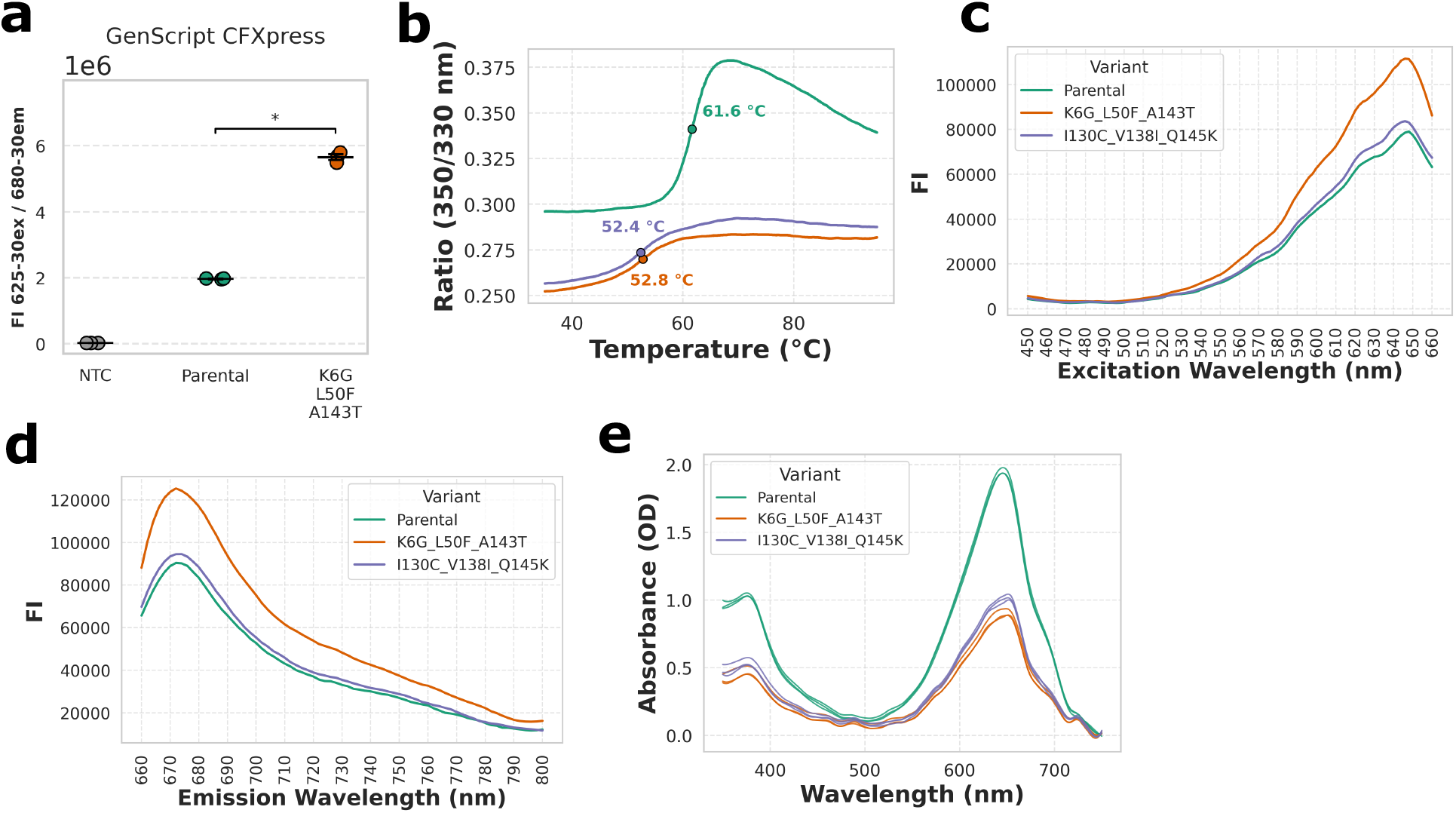
Purified protein characterization of top miRFP670nano3 variants. (a) Brightness of parental miRFP670nano3 and K6G-L50F-A143T variant in an alternate bacterial lysate based CFPS system. (b) Differential scanning fluorimetry profiles fitting inflection temperatures (T_i_). Proteins have not been normalized for concentration. (c) Excitation spectra (blank-corrected fluorescence, 450–660 nm). (d) Emission spectra (blank-corrected fluorescence, 660–800 nm). (e) Absorbance spectra (350–750 nm) of purified proteins used for quality control and concentration determination.

**Figure S4:**
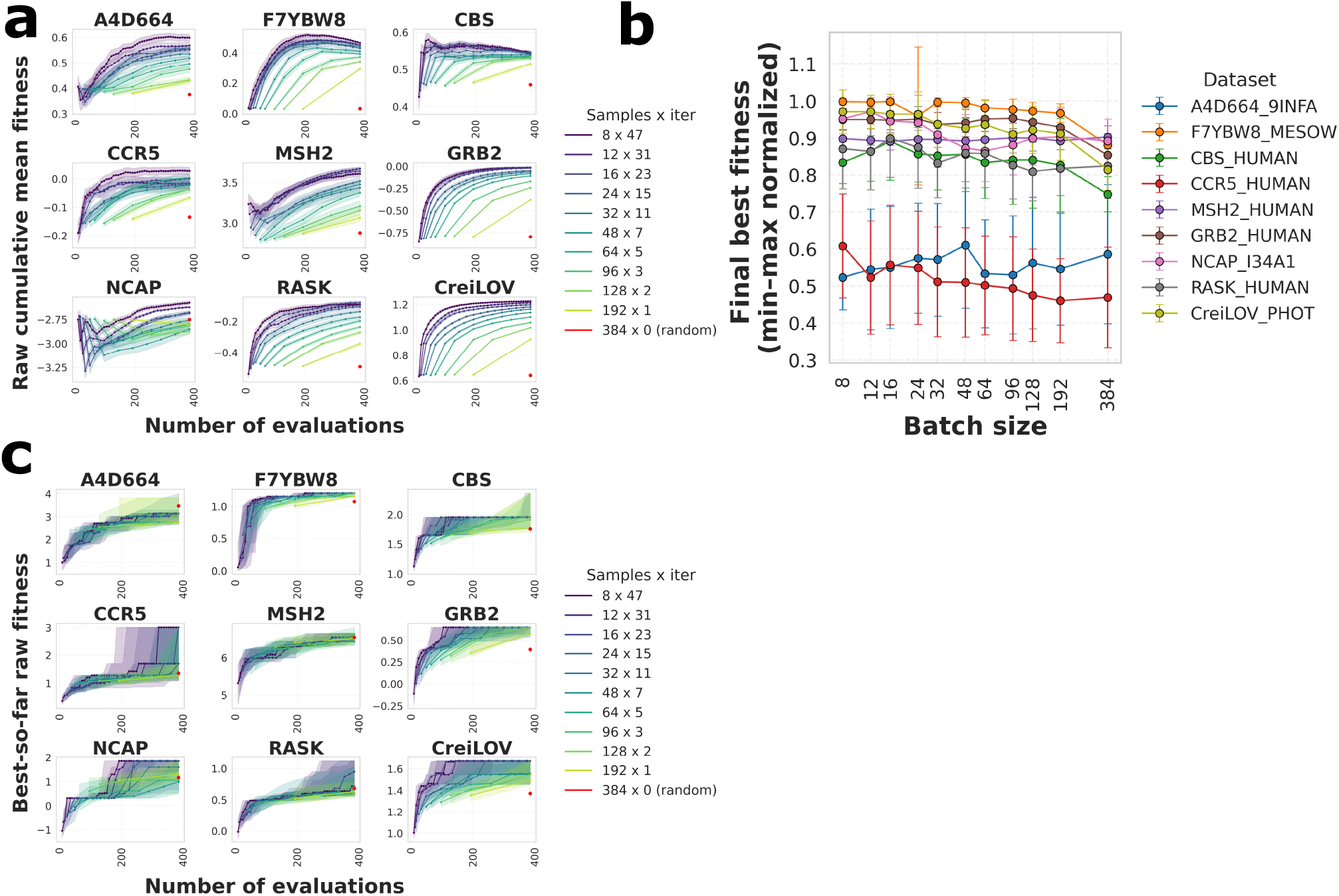
Additional batch-size benchmarking analysis. (a) Final best-fitness summary across acquisition batch sizes for each of the nine benchmark datasets. Error bars/shading represent sd of 20 random seeds. (b) Per-dataset convergence curves to the best variant found (best-so-far) across acquisition batch sizes. (c) Per-dataset convergence curves to cumulative-mean fitness across acquisition batch sizes.

**Figure S5:**
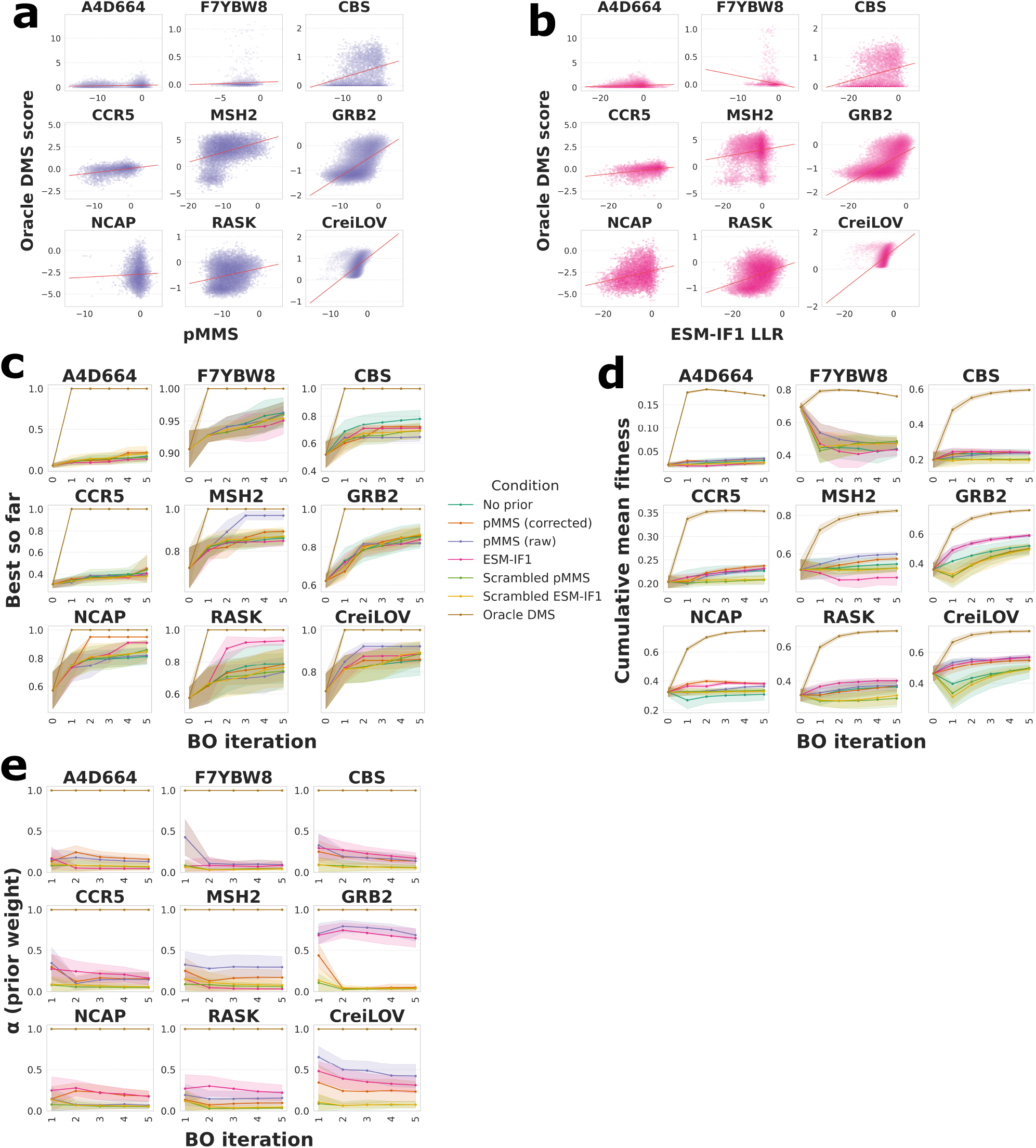
Additional information about GP prior-mean benchmarking. (a) Raw (uncorrected) masked-marginal score (pMMS) as zero-shot fitness score versus oracle DMS fitness from ProteinGym datasets. Pearson line of best fit is indicated. (b) As for a, with ESM-IF1 inverse-folding log-likelihood as the zero-shot fitness score. (c) Per-dataset convergence curves to the best variant found (best-so-far) across GP prior-mean conditions, including both raw and corrected pMMS. Corrected pMMS corrects for systematic substitution-identity bias (see Supporting Methods). (d) Per-dataset convergence curves to cumulative-mean fitness across GP prior-mean conditions, including both raw and corrected pMMS. (e) Per-dataset prior weight (*α*) across GP prior-mean conditions.

## Supporting Methods

### Cell-free protein expression

Parental miRFP670nano3 and its variants were ordered as linear synthetic genes (GBlocks) from Integrated DNA Technologies. Cell-free protein expression was conducted using the PURExpress In Vitro Protein Synthesis Kit (New England Biolabs E6800L). Reactions were conducted in a total volume of 2.5 µL. Templates were pipetted with F.A.S.T (Formulatrix) and master mixes with FlexDrop (Revvity). Reactions were carried out in triplicate with 250 pM template DNA along with 25 pM mNeonGreen DNA as a normalization control. Biliverdin was added to a final concentration of 5 µM, RNase inhibitor 0.1% (v/v) and samples were incubated at 37°C for 90 minutes.

For miRFP670nano3, fluorescence was measured with excitation at 625 nm (30 nm bandwidth) and emission at 680 nm (30 nm bandwidth). For mNeonGreen, excitation was at 488 nm (20 nm bandwidth) and emission at 535 nm (20 nm bandwidth).

Protein expression was quantified using the Nano-Glo HiBiT Lytic Detection System (Promega, N3030). LgBiT Protein was diluted 1:100, and Nano-Glo HiBiT Lytic Substrate was diluted 1:50 in Lytic Buffer. A volume of 2.5 µL of this mixture was added directly to the CFPS reaction with the FlexDrop. After a 10-minute room temperature incubation, luminescence was measured on a ClarioStar plate reader (BMG Labtech) at 470 nm emission with an 80 nm bandwidth.

Protein brightness and expression measurements from the laboratory were first normalized by dividing each assay measurement by the parental sequence mean. A single scalar activity label per variant was formed as a weighted combination of assays: brightness 75% and expression 25%. Per-sample means were taken over triplicate measurements and the resulting weighted activity values were used as training labels.

### Bayesian optimization configuration

A Gaussian process with a radial basis function kernel was then trained by exact marginal log likelihood, and an Upper Confidence Bound acquisition function (*β* = 1) selected 24 variants for each subsequent round. After three rounds of single mutants, a new design space was constructed comprising all three-way combinations of single mutants whose brightness or expression exceeded that of parental miRFP670nano3, and two further rounds were run identically. Each variant was tested for deviation from parental miRFP670nano3 with a one-sample t-test using a pooled variance estimate, with Benjamini-Hochberg correction.

### Active-learning directed evolution statistics

Each mutant was tested for significant deviation from wild-type using a one-sample t-test with a pooled vari-ance estimate. A one-sample test was performed due to the per-batch normalization of variants to parental miRFP670nano3. Pooled sample variance was derived from the within-group mean squared error (MSE) across each experimental batch. This approach is valid under the assumption of homoscedasticity across mutants, which is reasonable given that all variants were assayed under identical CFPS and plate-reader conditions. P-values were adjusted for multiple comparisons using the Benjamini-Hochberg method.

### Interpreting the Gaussian process model

GP performance was evaluated on the single-mutant subset (n = 72) using repeated 5-fold out-of-fold cross-validation. Samples were split into five folds with shuffling, and the procedure was repeated with 20 random seeds. Within each seed, samples were stratified by quantile bins of the target value (up to six bins, chosen so every bin held at least five members), falling back to a plain shuffled 5-fold split when no such binning was found. Folds were not grouped by residue position. Within each fold a StandardScaler was fit on the training data partition only and applied to the held-out fold; a GP was then fit on the resulting standardized ESM3 embeddings and predictions were generated for the held-out fold.

Evolutionary conservation of the mutations nominated during the miRFP670nano3 ALDE campaign was assessed against natural GAF-domain homologs. The top 25 mutations by weighted activity, spanning 20 unique positions (five positions contributed two nominated substitutions each), were analysed. The 147-aa GAF domain of miRFP670nano3 was used to query UniProt/NCBI by BLAST (E-value threshold 1× 10^*−*9^), and the BLAST-aligned subject sequences of the 1000 GAF-domain hits were globally realigned with MAFFT to give a consistent multiple sequence alignment. Each mutated position was mapped to its alignment column by counting non-gap residues in a reference aligned sequence, and Shannon entropy (excluding gaps) together with WT and mutant residue frequencies were computed per column. Substitutions were classified as “natural” versus “novel” using a 1% observed-frequency threshold for the mutant residue among homologs, within a Wilson-score 95% confidence interval. Shannon entropy at mutated positions was compared between the natural and novel groups with a Mann-Whitney U test (with rank-biserial correlation as effect size).

### Protein expression and purification

Plasmids in pET-28a backbone for bacterial expression were ordered from Genscript with C-terminal Flag and His tags. All constructs were validated by whole plasmid Oxford Nanopore sequencing (WEHI Genomics). Plasmids were co-transformed with Heme oxygenase 1 from *Synechocystis* (strain ATCC 27184) in pACYCDuet-1 in BL21(DE3) *E. coli* cells (New England BioLabs) using a heat-shock cycle. A single colony was picked to start a culture supplemented with kanamycin and chloramphenicol at 37°C. Protein expression was induced at optical density 0.6 - 0.8 by addition of 1 mM isopropyl *β*-D-1-thiogalactopyranoside (IPTG). Cells were pelleted after 16 h at 18°C and stored at -80°C until use.

Cells were thawed and resuspended with lysis buffer at 4°C for 30 minutes and then lysed via sonication on ice to ensure minimal thermal degradation. The resulting clarified lysate was incubated with 1 mL cOmplete His-Tag Purification resin (Roche) at 4°C for 2 hours. The solution was then passed through a gravity column and washed with DPBS + 50 mM Imidazole pH 8.0. Bound proteins were eluted with DPBS + 500 mM Imidazole pH 8.0. Eluted proteins were buffer exchanged into 20 mM Tris-HCl pH 8.0, 300 mM NaCl, 50 µM EDTA, 10% glycerol, 1 mM DTT, 0.01% NP-40 and concentrated. Protein concentration was quantified using a NanoDrop One/OneC spectrophotometer (ThermoFisher Scientific). Proteins were separated into aliquots and snap frozen using liquid nitrogen to store at -80°C.

### Biochemical characterization of miRFP670nano3 variants

Structural integrity and the inflection temperature (Ti) were tested using a Tycho NT.6 (Nanotemper) by measuring changes of the protein’s intrinsic fluorescence with the application of thermal ramp. Each protein was run as a single capillary with no technical replicates, and protein concentration was not normalized across capillaries prior to loading. A 200 µL aliquot of each sample recovered from the affinity chromatography step was injected onto a Superdex Increase 75 pg 10/300 GL column (Cytiva) for size exclusion chromatography (SEC) analysis. The column was equilibrated with DPBS and operated at 4°C. Fluorescence spectral properties were characterized using a CLARIOstar plate reader (BMG Labtech) in spectral-scan mode. For emission spectrum acquisition, samples were excited at 640 nm (10 nm bandwidth) and emission was collected from 665 to 801 nm in 2 nm steps (8 nm bandwidth). For excitation spectrum acquisition, emission was monitored at 690 nm (20 nm bandwidth) while the excitation wavelength was scanned from 450 to 660 nm in 2 nm steps (8 nm bandwidth). We derived Q-band to Soret-band absorbance ratios of ∼1.9 in our experiments, compared to a reference value of 3.2 for fully biliverdin-loaded protein. ^9^ We therefore report relative extinction coefficients and quantum yield values. All expression and purification steps were performed at the same time for all variants.

### Biological validation experiments in mammalian cell lines

HEK293T cells were validated by STR typing (AGRF, Australia) and confirmed to be mycoplasma negative by MycoAlert Kit (Lonza, LT07-118). 80,000 cells were seeded in 24-well plates (Corning, 3524) and transfected with 500 ng of plasmid DNA (pLenti-CMV-C-GFP-2A-Puro, Gen-script) per well using Lipofectamine 3000 (ThermoFisher, L3000015) at a 3:1 lipid-to-DNA ratio, following the manufacturer’s instructions. Cells were dissociated with 0.5% Trypsin-EDTA (Thermo 15400054) and analyzed on a NovoCyte Penteon Flow Cytometer (Agilent). Experiments were performed in biological triplicate

Stable miRFP670nano3-expressing lines were generated by lentiviral transduction. Lentivirus was produced via transfection of HEK293T cells using previously published methods. ^23^ Transduction of a fresh preparation of HEK293T cells was performed with 2*×* 10^5^ cells plated out the day before in 6-well plates. 2 mL fresh lentivirus supernatant supplemented with Polybrene 10 µg/mL (Sigma-Aldrich), was added to cells on the day of transduction. Fresh media was replaced on transduced cells 24 hours following infection. Parental and K6G-L50F-A143T miRFP670nano3 sequences were compared with and without an N-terminal LifeAct fusion, each across a serial dilution of viral supernatant, and far-red fluorescence was measured by flow cytometry as for transient transfection experiments. log_2_(geoMFI) was modeled as a quadratic function of log_2_(dilution) with a LifeAct (plus/minus) interaction on the linear term. Slope contrasts (differences in fitted log_2_(dilution) slopes between the four constructs) used standard errors propagated via the delta method from the model’s coefficient covariance matrix, with BH correction across the 6 pairwise comparisons.

### Retrospective simulation benchmarking

To guide ALDE design choices, we ran retrospective Bayesian optimization simulations on nine deep mutational scanning (DMS) landscapes from ProteinGym (Table 3), chosen to span a range of landscape shapes and difficulty.

**Table 3:** ProteinGym deep mutational scanning datasets used for retrospective ALDE simulation benchmarking.

| ProteinGym dataset ID |  |
| --- | --- |
| A4D664_9INFA | F7YBW8_MESOW |
| CBS_HUMAN | CCR5_HUMAN |
| MSH2_HUMAN | GRB2_HUMAN |
| NCAP_I34A1 | RASK_HUMAN |
| CreiLOV_PHOT |  |

To compare surrogate models, we simulated five rounds of Bayesian optimization (batch size 24, initial design of 24, 20 seeds) against a held-out pool of 384 variants, comparing the Gaussian process surrogate against ensemble baselines (an EvolvePro random forest and an ALDE XGBoost model with UCB acquisition) ^7,8^ and a random-sampling baseline. Performance was summarized per dataset (mean over seeds) and compared across the nine landscapes with a Friedman omnibus test and post-hoc Wilcoxon signed-rank tests of each active-learning method against the random baseline.

To evaluate the effect of varying the number of experimental batches and samples per batch, we swept various batch sizes and number of iterations for a fixed budget of 384 total evaluations, including a single batch of 384 sequences equivalent to random sampling. To test whether a zero-shot prior mean improves BO convergence, we compared seven GP prior-mean conditions across the nine datasets (20 seeds), holding the encoder, kernel and initialization fixed per seed so that only the prior differed. Scrambled (permuted) versions of each zero-shot score served as negative controls. The ground-truth DMS score acted as a positive control.

Raw pMMS is the masked-marginal log-odds fitness effect, summed over mutated positions and averaged per mutation: ^21^

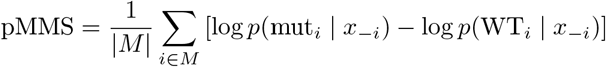

where *M* is the set of mutated positions in the variant, |*M* | its cardinality, *x*_*−i*_ the sequence context with position *i* masked, mut_*i*_ the mutant residue at position *i*, and WT_*i*_ the wild-type residue at position *i*; *p*(*·* |*x*_*−i*_) is the masked-language-model’s predicted probability of a residue at position *i* given that context.

However this raw score is systematically biased by substitution identity. ^22^ To adjust for this, we corrected pMMS z-scores each variant’s raw score against the mean and standard deviation of all single mutants sharing the same wild-type*→* mutant substitution type (minimum group size of five variants), following the substitution-type z-scoring approach of. ^22^ There was no significant difference in performance of raw and corrected pMMS across the nine datasets evaluated in our study (two-sided paired Wilcoxon signed-rank tests).

Prior informativeness was quantified as the Spearman correlation between the prior score and oracle DMS fitness. Benefit was quantified as the per-seed fractional regret reduction in the normalized area under the cumulative-mean curve relative to the no-prior baseline. For the CreiLOV pMMS/ESM-IF1 comparison, both priors outperformed no-prior in all 20 of 20 seeds. Because the sample size is small (*n* = 20), we used the exact Wilcoxon signed-rank test rather than the normal approximation, which can otherwise report *p*-values below what the test can actually resolve. The exact test returned its minimum attainable *p*-value (2*/*2^20^ = 1.9 *×*10^*−*6^), reflecting the fully saturated result (20/20 seeds favoring the prior).

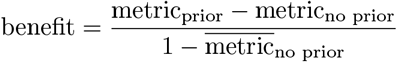

### Software versions

Analyses used Python 3.11 with PyTorch 2.5.1. Protein language model embeddings were generated with ESM3. Multiple sequence alignment used MAFFT v7.525.ΔΔG stability predictions used FoldX v5.1. ^17^ Gaussian process surrogates and Bayesian optimization used GPyTorch v1.13 and BoTorch v0.12.0. Feature standardization used scikit-learn v1.8.0. The ALDE XGBoost benchmark baseline used xgboost-distribution v0.4.0. The EvolvePro benchmark baseline followed the method of, ^8^ reimplemented with scikit-learn’s `RandomForestRegressor`.

## Notes

https://github.com/WEHIGenomicsRnD/Manuscript_FRP_MLDE

https://github.com/WEHIGenomicsRnD/MLDE_qHSRI

### Competing Interest Statement

The authors have declared no competing interest.

